# Engineering an *fgfr4* knockout zebrafish to study its role in development and disease

**DOI:** 10.1101/2024.05.08.593184

**Authors:** Emma N. Harrison, Amanda N. Jay, Matthew R. Kent, Talia P. Sukienik, Collette A. LaVigne, Genevieve C. Kendall

## Abstract

Fibroblast growth factor receptor 4 (FGFR4) has a role in many biological processes, including lipid metabolism, tissue repair, and vertebrate development. In recent years, FGFR4 overexpression and activating mutations have been associated with numerous adult and pediatric cancers. As such, *FGFR4* presents an opportunity for therapeutic targeting which is being pursued in clinical trials. To understand the role of FGFR4 signaling in disease and development, we generated and characterized three alleles of *fgfr4* knockout zebrafish strains using CRISPR/Cas9. To generate *fgfr4* knockout crispants, we injected single-cell wildtype zebrafish embryos with *fgfr4* targeting guide RNA and Cas9 proteins, identified adult founders, and outcrossed to wildtype zebrafish to create an F1 generation. The generated mutations introduce a stop codon within the second Ig-like domain of Fgfr4, resulting in a truncated 215, 223, or 228 amino acid Fgfr4 protein compared to 922 amino acids in the full-length protein. All mutant strains exhibited significantly decreased *fgfr4* mRNA expression during development, providing evidence for successful knockout of *fgfr4* in mutant zebrafish. We found that, consistent with other *Fgfr4* knockout animal models, the *fgfr4* mutant fish developed normally; however, homozygous *fgfr4* mutant zebrafish were significantly smaller than wildtype fish at three months post fertilization. These *fgfr4* knockout zebrafish lines are a valuable tool to study the role of FGFR4 in vertebrate development and its viability as a potential therapeutic target in pediatric and adult cancers, as well as other diseases.

## Introduction

Fibroblast growth factor 4 (FGFR4) is a member of the fibroblast growth factor receptor family, a group of receptor tyrosine kinases that bind promiscuously to fibroblast growth factors (FGFs) and play a role in a wide range of biological processes. FGFR family members regulate a wide-range of processes including organogenesis, neural development, metabolism, and inflammation (1). FGFR4 signaling is critical in chordate development, tissue repair, and lipid metabolism. Additionally, FGFR4 has been implicated as a crucial mediator of myogenesis and muscle regeneration in mouse and chick models (2, 3).

In recent years, FGFR4 has been implicated as a potential therapeutic target in rhabdomyosarcoma (RMS), a pediatric soft tissue sarcoma. RMS is most often associated with displaying molecular features of the skeletal muscle lineage and is characterized by its failure to terminally differentiate into mature muscle (4, 5). Though caused by multiple distinct genetic drivers, the subtype in which FGFR4 has been most often implicated is fusion-positive rhabdomyosarcoma. This subtype also has poorer patient prognosis compared to fusion-negative RMS subtypes (6, 7). Fusion-positive RMS is most often genetically driven by a single chimeric fusion oncogene *PAX3/7::FOXO1* which is caused by a chromosomal translocation event wherein the DNA binding domain of *PAX3/PAX7* is juxtaposed to the transactivation domain of *FOXO1* (8-10). FGFR4 is transcriptionally upregulated in fusion-positive RMS, and its overexpression is a predictor for reduced overall survival in patients (11, 12). FGFR4 has also been established as a direct transcriptional target of the PAX3::FOXO1 fusion protein (13, 14).

Beyond rhabdomyosarcoma, FGFR4 has also garnered attention in recent decades for its dysregulation in other adult and pediatric cancers. FGFR4 is frequently overexpressed in cancers, and it is prone to hotspot activating mutations in breast cancer and hepatocellular carcinoma, among others (11, 15). FGFR4 is an exciting therapeutic target because many pan-FGFR inhibitors and FGFR4 inhibitors have been developed, three of which are currently FDA approved for cancer treatment (16). Multiple phase 1 and phase 2 clinical trials are currently underway using select small molecule inhibitors of FGFR4 in cancers such as hepatocellular carcinoma and acute myeloid leukemia (17). In addition, CAR T-cell therapy targeting FGFR4 in rhabdomyosarcoma is in preclinical development stages and has had success in mouse allograft models to decrease tumor burden (18).

Though broadly regarded as a therapeutic target of interest, a mechanistic understanding of the role of FGFR4 in rhabdomyosarcoma and other cancers remains unclear. Here, we have developed and thoroughly characterized three alleles of *fgfr4* knockout zebrafish to fully understand the phenotypic consequences of the loss of this gene. These knockout zebrafish lines can be used to study the role of *fgfr4* in biological processes related to development and disease. This knockout model can also be utilized in existing zebrafish disease models to thoroughly understand the role of FGFR4 in various diseases, including rhabdomyosarcoma and other cancers.

## Methods

### Zebrafish husbandry and breeding

*Danio rerio* were housed in an AAALAC-accredited, USDA-registered, OLAW-assured Aquaneering facility in compliance with the Guide for the Care and Use of Laboratory Animals. Vertebrate animal work was overseen by the Abigail Wexner Research Institute at Nationwide Children’s Hospital IACUC committee and Animal Resources Core. WIK zebrafish (Zebrafish International Resource Center, ZL84) served as our wildtype fish. Each strain was separately reared to adulthood (∼3 months in our facility) and bred in-house by mass breeding. Larval fish (5-30 days post-fertilization) were housed at a density of approximately 15 larvae/L. Adult fish were housed at a density of 5-10 fish/L in mixed-sex groups. Fish were housed in 2.8L and 6L tanks on a recirculating life support system (Aquaneering, San Diego, CA) supplied with reverse osmosis water. An exchange of 10-15% of the total system water volume with fresh reverse osmosis water occurs daily. Fgfr4 mutant lines were outcrossed every two generations to maintain genetic diversity. The system water was maintained at 26-27.5 °C, conductivity of 450– 600 μS/cm, and a pH of 7.2-7.5 using continuous monitoring via probes in the sump and an automatic dosing system. Larval fish were fed three times a day with 30,000 L-type rotifers *Brachionus plicatilus* gut loaded with RGComplete rotifer feed (Reed Mariculture) per 1.8L tank. Adult fish were fed twice a day with Gemma Micro 300 (Skretting).

### Generating fgfr4 knockout fish

CRISPR/Cas9 knockout adapted from Talbot and Amacher, 2014 (19). *fgfr4* gRNA (IDT, see **Supplementary Table 1** for primer sequences) and Cas9 protein (IDT, 1081060) were injected into the single cell of WIK embryos to generate potential founders. Once sexually mature, potential founders were outcrossed to wildtype zebrafish to separate mosaic CRISPR-mediated mutations. After three *fgfr4* mutant strains were identified by sequencing, heterozygous fish of each strain were in-crossed to generate homozygote zebrafish colonies.

### High-Resolution Melt Analysis (HRMA)

HRMA was performed as described previously (20). Genomic DNA was isolated from the caudal fins of potential mutant zebrafish, as well as a minimum of three wildtype (WIK) fins as a reference group. Fin clips were incubated in 10mM Tris HCl (Sigma, 10708976001) pH 8.3, 50mM KCl (Sigma, P5405-250G), 0.3% Triton X-100 (Fisher, BP151-100), and 0.3% NP40 (Fisher, NC9375914) at 98°C for 10 minutes, then cooled to 4°C for 10 minutes. One-tenth volume of 10mg/mL proteinase K (Fisher, BP1700-100) was added to samples, which were then incubated for at least 1.5 hours at 55°C, proteinase K heat inactivated at 95°C for 10 minutes and held at 4°C until ready for use. Genomic DNA was added to 9 μL of Precision Melt Supermix (Bio-Rad, BP1700-100) and 0.5 μL of each 10 μM forward and reverse primers targeting the mutated region in *fgfr4* (IDT, see **Supplementary Table 1** for primer sequences) and diluted to 20 μL with nuclease free water in a 384-well plate (Bio-Rad, HSP3805). The plate was run on a Bio-Rad CFX384 Real Time PCR Detection System with the following program: 95 °C for 3 min; (95 °C for 15 s, 60 °C for 20 s, 70 °C for 20 s) x 45 cycles; 65 °C for 30 s; melt 65°C–95 °C, 0.2 °C/step hold 5 s; 95 °C for 15 s. Bio-Rad Maestro and Bio-Rad Precision Melt Analysis software were used for data analysis, with known wildtype samples as the reference group.

### A-Tail Cloning

To identify the sequence of CRISPR-mediated mutations in the *fgfr4* gene, genomic DNA used in the HRMA was used in a Phusion based PCR with 10 μM forward and reverse primers (see **Supplementary Table 1** for primer sequences). PCR products were purified using the Monarch PCR Cleanup Kit (NEB, T1030L). PCR products were A-tailed, ligated into the PGEM-T Easy vector (Promega, A1360), and then transformed into DH5-α cells (Fisher Scientific, FEREC0111). DH5-α cells were plated on LB-Amp plates sprayed with X-Gal/IPTG (Fisher Scientific, 21530077) and incubated overnight at 37°C. Colonies were picked, cultured overnight, and mini-prepped with QIAGEN QIAprep Spin Miniprep Kit (27106), and DNA was sequenced by Sanger sequencing through Eurofins Genomics using the SP6 sequencing primer. For sequence alignment, the wildtype reference used was *Danio rerio* fgfr4 cDNA from genome assembly GRCz11, NM_131430.1, chr21:37183912-37194363.

### Genotyping

Zebrafish were anesthetized in 0.2 mg/mL tricaine methanesulfonate (Fisher Scientific, NC0342409) and the tail fin was cut. Fin clips were incubated in 10mM Tris HCL pH 8.3 (Sigma, 10708976001) pH 8.3, 50mM KCl (Sigma, P5405-250G), 0.3% Triton X-100 (Fisher, BP151-100), and 0.3% NP40 (Fisher, NC9375914) at 98°C for 10 minutes, then cooled to 4°C for 10 minutes. One-tenth volume of 10mg/mL proteinase K (Fisher, BP1700-100) was added to samples, which were then incubated overnight at 55°C, proteinase K heat inactivated at 95°C for 10 minutes, and held at 4°C. The PCR reaction mix was created with NEB Phusion GC Buffer (NEB, B0519S), Phusion DNA Polymerase (NEB, M0530S), 10mM dNTPs (Fisher Scientific, 10-297-018), 10 μM forward primer and 10 μM reverse primer (IDT) according to the recommended ratios (see **Supplementary Table 1** for primer sequences). The samples were then amplified using the following program: 98°C x 30 sec, 98°C x 10 sec, 60°C x 30 sec, 72°C x 30 sec, repeat the previous three steps for 34 cycles (35 total cycles), 72°C x 10 min, and HOLD at 12°C. PCR products were purified using the Monarch PCR Cleanup Kit (NEB, T1030L) and digested with the *TfiI* restriction enzyme (NEB, R0546S) digest. Final products were run in a 1.2% agarose gel (Fisher Scientific, BP160-500) and imaged on a Bio-Rad Molecular Imager Gel Doc XR+ with Image Lab Software. With this genotyping method, the expected product size(s) for a wildtype zebrafish are 89 and 72 base pairs (bp), for a heterozygous fish are 163, 89, and 72bp, and for a homozygous fish is 163bp.

### qRT-PCR

Wildtype WIK and fgfr4 homozygous mutant adult zebrafish were set up in breeding chambers with dividers and left overnight. The following morning, dividers were pulled, and embryos were collected. At precisely 24 hours post fertilization, embryos of each cross were dechorionated either manually or via pronase (Sigma-Aldrich, 11459643001) dechorionation, euthanized, and aliquoted into samples of 12 embryos each. Extra embryo media was eluted off the embryo pellet, and samples were snap frozen and stored at -80°C.

Total RNA was isolated from each sample using the QIAGEN RNeasy Mini Kit (74104) including the optional on-column DNAse digestion. The RT2 HT First Strand Synthesis kit (Qiagen, 330411) was used to synthesize cDNA using approximately 825 ng of total RNA input per sample. cDNA for each sample was diluted to 80 μL with nuclease-free water. 4 μL of diluted cDNA, 5 μL Universal SYBR Green Supermix (Bio-Rad, 1725122), and 0.5 μL of forward and reverse primers (see **Supplementary Table 1** for primer sequences) for the genes of interest were combined in a 384-well plate (Bio-Rad, HSP3805). The plate was run on a CFX384 Real Time PCR Detection System using the following program: 95 °C for 2 min; (95 °C for 15 s, 60 °C for 1 min, Plate Read) x 40 cycles; 65 °C for 30 s; (65 °C, 0.5 °C/step, Plate Read) x 60 cycles. Five biological replicates were used per strain with three technical replicates each. Data was analyzed using the CFX Maestro program.

### Histology

Male and female adult homozygous mutant zebrafish from each strain were euthanized, placed in a cassette, and fixed in 4% PFA (Fisher Scientific, 50-276-31) in PBS for a minimum of 24 hours at room temperature. Cassettes were then transferred into 0.5M EDTA pH 7.8 (Fisher Scientific, BP120-1) for five days, after which zebrafish were paraffin embedded, sagittally sectioned, and hematoxylin and eosin (H&E) stained as described previously (21).

### Imaging of zebrafish embryos and adults

Images of 24 hours post-fertilization (hpf) and 48hpf zebrafish embryos were taken on a Leica M205FA dissecting microscope. Images of adult zebrafish were taken by anesthetizing zebrafish in 0.2 mg/mL tricaine methanesulfonate (MS-222), placing them on laminated grid paper, and taking images with a smartphone camera. Standard length was measured from anterior-most part of the head to the posterior-most part of the tail in embryos, or to the caudal fork of the tail fin in adults.

### Quantification and statistical analysis

All experimental statistical tests were performed in Graphpad Prism 9 (La Jolla, CA). Sample sizes and the statistical tests performed are provided in figures or figure legends.

## Results

Knockout *fgfr4* zebrafish were generated using a CRISPR/Cas9 strategy (**Fig 1A**). Guide RNA against zebrafish *fgfr4* and Cas9 protein were injected into single-cell wildtype embryos. After outcrossing these founders to wildtype to isolate individual CRISPR-mediated germline mutations, genomic DNA from putative F1 mutant fish was used for high resolution melt analysis (HRMA) to identify differences in melting temperature compared to wildtype references, which is indicative of genetic mutations. HRM analysis identified three clusters with inflection curves distinct from the wildtype references (**Fig 1B**). Sub-cloning and Sanger sequencing revealed that each of the clusters possessed one of the following mutations in the *fgfr4* locus when aligned to wildtype *fgfr4* cDNA: a 7 base pair (bp) deletion at 604-610bp (GATTCAG/-) and 6bp deletion at 644-649bp (AGGCTC/-), two 1bp insertion after 605bp (-/G) and 606bp (-/A), one substitution at 609bp (A/T), 38bp deletion at 609-646bp (CAGGAGTATATGTGTGTATGCTACGTGGCACCAAAGAG/-), and a 1bp insertion after 669bp (-/G) (**Fig S1**). Each of these genomic mutations conferred premature stop codons in the *fgfr4* sequence, resulting in truncated protein sequences with predicted lengths of 223, 228, and 215 amino acids (**Fig 1C**). These alleles are referred to as fgfr4^nch4^, fgfr4^nch5^, and fgfr4^nch6^, respectively. Because the kinase domain of fgfr4 is lost in the generated mutants, all downstream wildtype function should also be lost.

**Figure 1.**
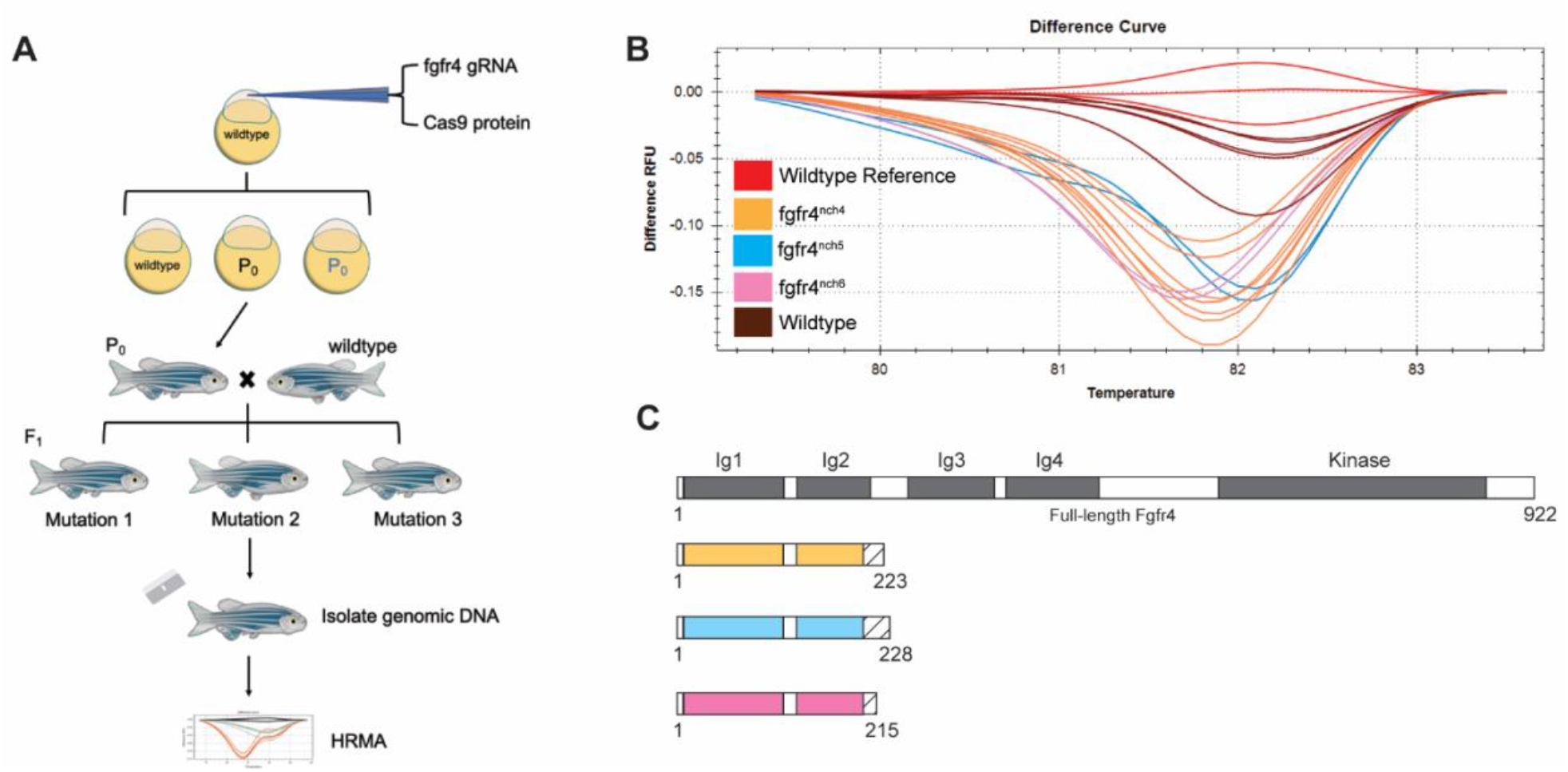
Three strains of zebrafish *fgfr4* knockout mutants generated using CRISPR/Cas9. (A) Schematic of generation of *fgfr4* knockout zebrafish. Single cell wildtype zebrafish embryos were injected with *fgfr4* guide RNA and Cas9 protein to generate mutant founders. Potential founders were outcrossed to isolate CRISPR-mediated mutations, and putative crispant F1s were subjected to high resolution melt analysis to identify mutation sequences. (B) High resolution melt analysis (HRMA) of potential F1 mutants revealed three distinct clusters of potential *fgfr4* crispants. (C) Schematic of wildtype Fgfr4 and predicted knockout strain protein lengths. Premature stop codons produced truncated Fgfr4 proteins with predicted length of 223, 228, and 215 amino acids respectively. Solid blocks indicate regions of wildtype Fgfr4 amino acid sequence alignment. The protein sequence aligned to the wildtype Fgfr4 protein ends at the same amino acid for all three alleles, with various additions of amino acids that do not align to the wildtype protein afterwards. Hatching indicates these additional amino acids.

We used quantitative real-time polymerase chain reaction (qRT-PCR) as a proxy to validate the *fgfr4* genomic knockout in all strains. To assess *fgfr4* mRNA abundance, we designed qRT-PCR primers to target regions both 5’ and 3’ of the mutation site in all strains (**Fig 2A**). In all putative knockout strains, relative expression of *fgfr4* mRNA was significantly decreased in homozygous *fgfr4* knockout embryos compared to wildtype embryos (**Fig 2B-C**), indicating that the *fgfr4* mRNA transcript is less stable, indicative of a true knockout. Because it has the lowest *fgfr4* mRNA expression of all strains, the fgfr4^nch4^ line was used for the remainder of the described experiments, though all other strains had consistent phenotypes with those described below (data not shown).

**Figure 2.**
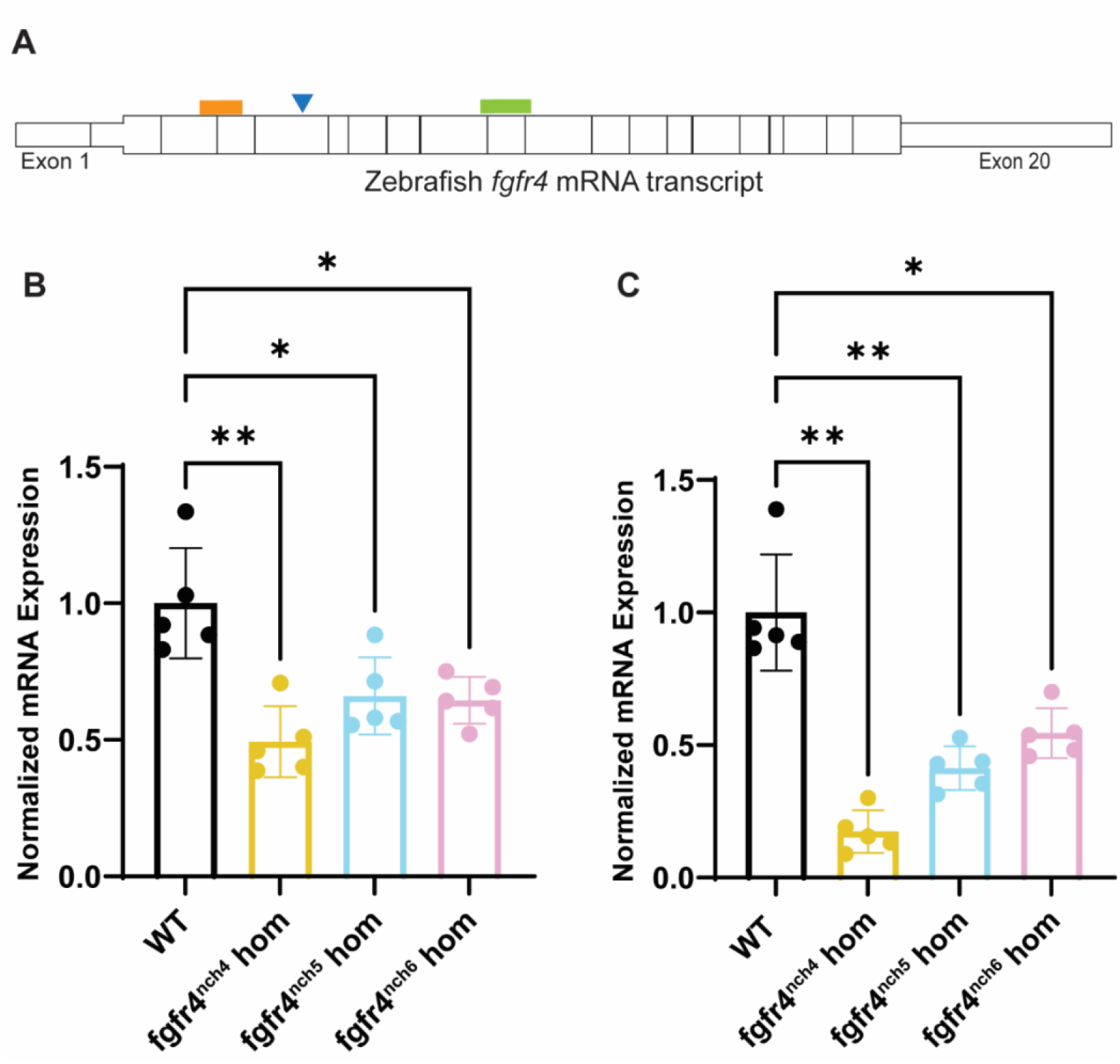
Zebrafish *fgfr4* knockout validated by quantitative reverse transcription polymerase chain reaction (qRT-PCR). (A) Schematic of wildtype zebrafish *fgfr4* mature mRNA transcript and regions targeted for qRT-PCR. Blue triangle indicates approximate *fgfr4* crispant mutation site for all strains relative to primers. Orange box indicates region 5’ of mutation site targeted by qRT-PCR primers, and green box indicates region 3’ of mutation site targeted by qRT-PCR primers. Relative expression of *fgfr4* mRNA as measured by qRT-PCR targeting the regions 5’ (B) and 3’ (C) of the sequence mutation. Expression is normalized to *gapdh* and *rpl13a*. Each point represents a pool of n=12 24 hours post-fertilization (hpf) zebrafish embryos derived from a maternal zygotic. Multiple points represent biological replicates, while three technical replicates were used to generate each biological replicate. Error bar is the mean ± standard deviation. P values were calculated using a one-way Brown-Forsythe and Welch ANOVA, correcting for multiple comparisons with a Dunnett T3 test.

Homozygous *fgfr4* mutant zebrafish embryos did not appear phenotypically different from wildtype zebrafish at early developmental timepoints (**Fig 3A**). Homozygous fgfr4 mutant zebrafish are not significantly different in size than stage-matched wildtype embryos at 24- and 48-hours post-fertilization (hpf). This standard early embryonic development is consistent with other zebrafish *fgfr4* knockout models (22). Further, all three fgfr4 mutant zebrafish alleles were found to exhibit Mendelian ratios (**Supplementary Table 2**), and homozygous *fgfr4* fish were viable and fertile. Sex distributions were also standard in fgfr4 homozygotes (**Supplementary Table 3**). However, a size discrepancy between *fgfr4* homozygotes and wildtype fish became apparent at roughly three months post-fertilization, wherein *fgfr4* homozygotes are significantly smaller than age-matched wildtype zebrafish (**Fig 3B**), though this size discrepancy becomes nonsignificant at six months post-fertilization (**Fig S2**). However, the internal anatomical histology of adult fgfr4 knockout zebrafish appeared overall normal (**Fig 3C**).

**Figure 3.**
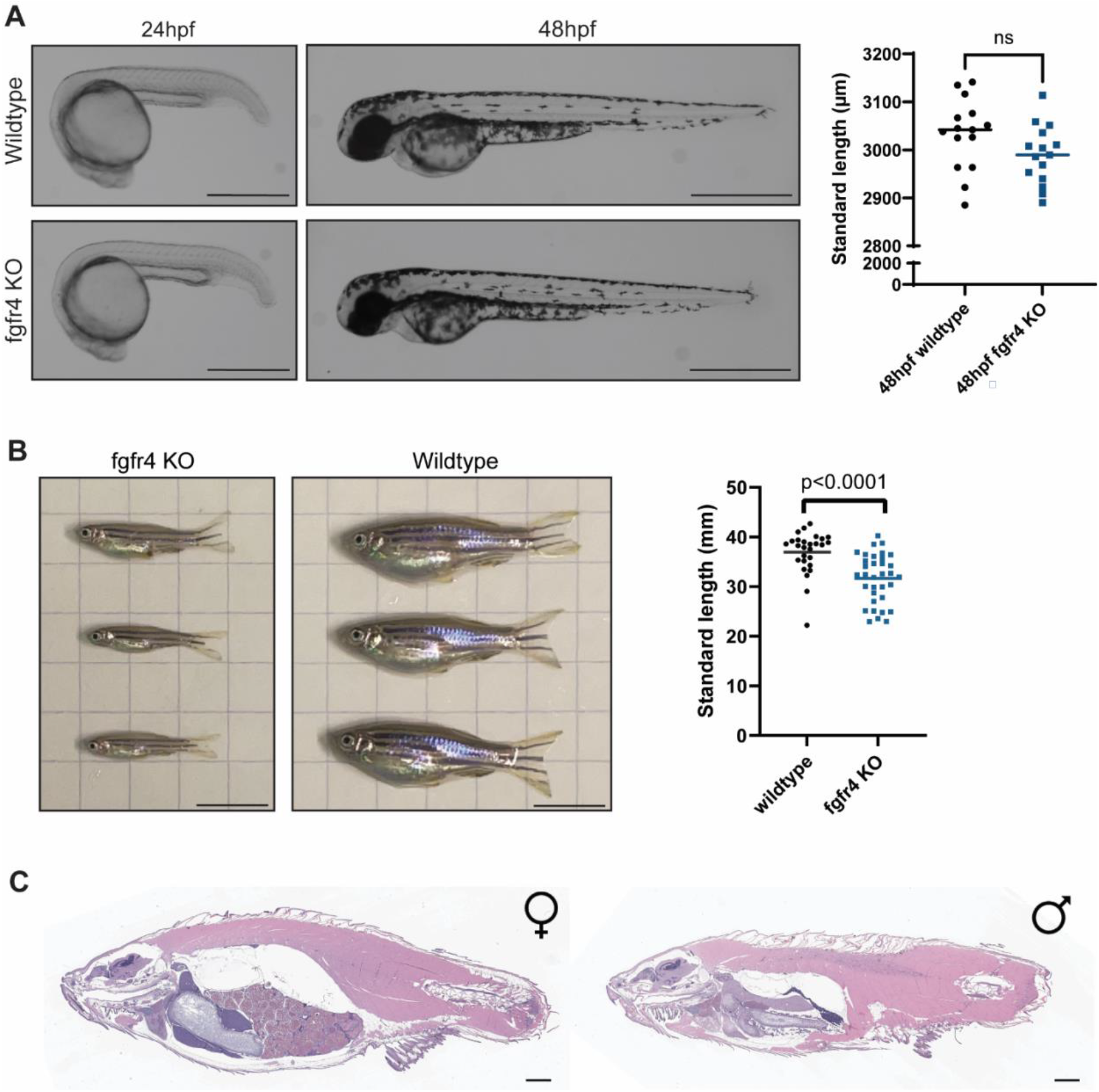
Homozygous *fgfr4* knockout (KO) zebrafish have no embryonic phenotype but are significantly smaller than wildtype zebrafish at three months post-fertilization (mpf). (A) Representative images from a phenotypic analysis of embryonic zebrafish. Homozygous *fgfr4* mutant zebrafish do not display an early embryonic phenotype. Dots represent individual fish standard lengths, and the bar represents the mean. An unpaired two-tailed t-test with Welch’s correction was used to calculate the p value. Scale bar is 560.1 μm in 24 hpf images and 700.4 μm in 48 hpf images. Standard length quantification was performed with n=15 embryos per group. (B) Phenotypic analysis of adult zebrafish. Homozygous knockout *fgfr4* zebrafish are significantly smaller than wildtype zebrafish at three months post fertilization. Standard length quantification was performed with n=27 WT fish and n=34 *fgfr4* KO adult fish. Each point represents an individual fish standard lengths, and the bar represents the mean. An unpaired two-tailed t-test with Welch’s correction was used to calculate p value. Scale bar is 1 cm. (C) Representative hematoxylin and eosin (H&E) staining of a sagittal section from female and male adult *fgfr4* knockout zebrafish. Internal anatomy in mutant zebrafish is normal in both male and female adult *fgfr4* knockout zebrafish. This was assessed in 5 female and 3 male *fgfr4* knockout fish total. Scale bar is 1mm.

## Discussion

Here, we developed three *fgfr4* knockout strains through a CRISPR/Cas9 single-cell injection strategy. Putative knockout strains were identified by HRMA, and the loss of *fgfr4* expression was validated by qRT-PCR. Because all mutations confer premature stop codons that eliminate the kinase domain of Fgfr4, normal Fgfr4 activity should be removed in all mutants. Fgfr4 is not duplicated in zebrafish indicating that this single knockout should remove all Fgfr4 activity. The *fgfr4* mutant strains were embryonically viable and were phenotypically normal at early developmental timepoints. However, at approximately three months post fertilization, juvenile *fgfr4* knockout zebrafish are significantly smaller than age-matched wildtype fish. Despite their relatively small size at this timepoint, mutant *fgfr4* zebrafish have normal internal anatomy and are fertile.

The results seen in this study recapitulate what has been seen in existing *fgfr4* animal models. Similarly to zebrafish, *Fgfr4* knockout mice are embryonically viable, develop normally, and experienced no impacts on fertility (23). *Fgfr4* knockout mice also metabolize high-fat diets differently than wildtype mice (24). An *fgfr4* zebrafish knockout model was similarly viable and developed normally in early stages of development (22), though adult phenotypes were not fully characterized. In zebrafish, the viability and lack of embryonic phenotype is likely a result of compensation from other FGFR family members (22).

The smaller size of *fgfr4* mutant zebrafish in the juvenile stage has also been noted in *Fgfr4* knockout mice, which weigh 10% less than their wildtype siblings at the time of weaning (23). This difference in size may be due to the established and conserved role of *fgfr4* in cholesterol metabolism. Because zebrafish embryos obtain all nutrition from the yolk until about five days post-fertilization, the metabolism-related phenotypic outcomes of an *fgfr4* knockout may not appear until larvae are able to feed freely.

In the future, we plan to use these knockout zebrafish to understand the genetic cooperation between FGFR4 and PAX3::FOXO1, the predominant driver of fusion-positive rhabdomyosarcoma. We have found that when human PAX3::FOXO1 is incorporated into the zebrafish genome, we obtain tumors that histologically and molecularly recapitulate human RMS (25). We hypothesize that incorporating PAX3::FOXO1 into the genome of fgfr4 knockout zebrafish, tumor occurrence and tumor volume will be significantly reduced. The *fgfr4* knockout zebrafish strains described here are a valuable tool in the study of the role of FGFR4 in vertebrate development and disease.

## Acknowledgements

We would like to acknowledge the Animal Resources Core at Nationwide Children’s Hospital for phenomenal caretaking of our zebrafish colonies, in particular Dr. Laurie Goodchild, Dr. Carmen Arsuaga, Dr. Lindsey Ferguson, Logan Fehrenbach, Logan Bern, and Alexander Kramer. We also thank the Histopathology Core and the Processing and Banking Core at Nationwide Children’s Hospital for the processing and imaging of our histology samples.

## Data Availability

All data generated in this study are included in this article and supplementary files. We are in the process of making all three fgfr4 knockout zebrafish lines available through the Zebrafish International Resource Center (ZIRC) for their use by the broader scientific community.

## Ethics Statement

*Danio rerio* were housed in an AAALAC-accredited, USDA-registered, OLAW-assured Aquaneering facility in compliance with the Guide for the Care and Use of Laboratory Animals. Vertebrate animal work was overseen by the Abigail Wexner Research Institute at Nationwide Children’s Hospital IACUC committee and Animal Resources Core.

## Author Contributions

G.C.K. conceived and supervised the study. E.N.H., A.N.J., M.R.K., T.P.S., and C.A.L. performed experiments. E.N.H., A.N.J., M.R.K, T.P.S, and C.A.L. analyzed experimental data. E.N.H., A.N.J., and G.C.K. drafted the manuscript and figures. All authors reviewed and edited the final manuscript. A.N.J., M.R.K., and G.C.K. obtained funding.

