## Supplementary Materials for "Engineering an *fgfr4* knockout zebrafish to study its role in development and disease"

| **Oligonucleotides** | **Sequence** |
|---|---|
| *fgfr4* gRNA | Guide 1: GAATCTTCATAGGTAACACCGA<br><br>Guide 2: AAGAGGCTCTACGCAACTCCGA |
| *fgfr4* HRMA/A-Tail Cloning primers | FWD: 5'-GTGGTTCAAGAGTGGTGTTC-3'<br><br>REV: 5'-CTGCCACTGTGATAGTGAAG-3' |
| *fgfr4* genotyping primers | FWD: 5'-CCAGTGGTTCAAGAGTGGTGTTC-3'<br><br>REV: 5'- CTACCTGCCACTGTGATAGTGAAG-3' |
| *fgfr4* qRT-PCR primers (5' of mutation) | FWD: 5'-AGTGGTTCAAGAAGGACAGTAA-3'<br><br>REV: 5'-AAGGAAATATGTTGACTTTGGAGAG-3' |
| *fgfr4* qRT-PCR primers (3' of mutation) | FWD: 5'-GTGAAGATTGTGAAGACAGGAAGC-3'<br><br>REV: 5'-GGCTACATCCTCCTCTGATAAGAC-3' |
| *gapdh* qRT-PCR primers (Kendall et al., 2018) | FWD: 5'-GTGGCCATCAATGACCCATTC-3'<br><br>REV: 5'-CAATGACCAGTTTGCCGCCTTC-3' |
| *rpl13a* qRT-PCR primers (Kendall et al., 2018) | FWD: 5'-CGGTCGTCTTTCCGCTATT-3'<br><br>REV: 5'-TTCCAGAGATGTTGATACCCTCAC-3' |

**Supplementary Table 1. Oligonucleotide sequences**

| | **fgfr4$^{nch4}$** | | **fgfr4$^{nch5}$** | | **fgfr4$^{nch6}$** | |
|---|---|---|---|---|---|---|
| | **Number** | **%** | **Number** | **%** | **Number** | **%** |
| **Homozygotes** | 35 | 17.12 | 38 | 21.69 | 26 | 28.4 |
| **Heterozygotes** | 57 | 51.35 | 110 | 58.2 | 32 | 39.5 |
| **Wildtype** | 19 | 31.53 | 41 | 20.11 | 23 | 32.1 |
| **Total** | 111 | 100 | 189 | 100 | 81 | 100 |

**Supplementary Table 2. Genotypes of *fgfr4* knockout strains**. Genotypes obtained from heterozygous in-crosses for each *fgfr4* allele. Differences between observed and expected genotypes were nonsignificant by chi-square statistical tests. P values for actual and expected genotypes were 0.2950, 0.2706, and 0.3917 for fgfr4$^{nch4}$, fgfr4$^{nch5}$, and fgfr4$^{nch6}$ respectively.

| | fgfr4$^{nch4}$ | | fgfr4$^{nch5}$ | | fgfr4$^{nch6}$ | |
|---|---|---|---|---|---|---|
| | Number | % | Number | % | Number | % |
| **Female** | 14 | 58.3% | 28 | 54.9% | 13 | 46.4% |
| **Male** | 10 | 41.7% | 23 | 45.1% | 15 | 53.6% |
| **Total** | 24 | 100.0% | 51 | 100.0% | 28 | 100.0% |

**Supplementary Table 3. Sex split of homozygous *fgfr4* knockout zebrafish.** Sex split data obtained from homozygous *fgfr4* mutant zebrafish colonies. Differences between observed and expected sex distributions were nonsignificant by chi-square statistical tests. P values for actual and expected sexes were 0.6184, 0.7891, and 0.5623 for fgfr4$^{nch4}$, fgfr4$^{nch5}$, and fgfr4$^{nch6}$ respectively.

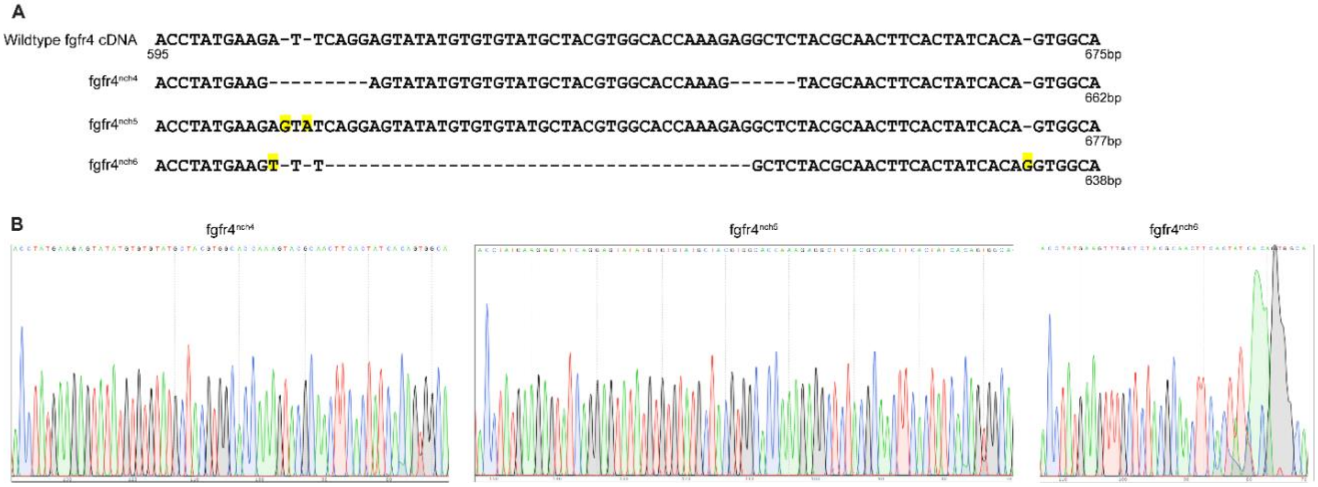


**Supplementary Figure 1. Mutation sequences identified by Sanger sequencing.** (A) Mutation sequences alignments to wildtype *fgfr4* cDNA. Wildtype reference is *Danio rerio* fgfr4 cDNA from genome assembly GRCz11, NM_131430.1, chr21:37183912-37194363. Strain fgfr4$^{nch4}$ contains a 7 base pair (bp) deletion at 604-610bp (GATTCAG/-) and 6bp deletion at 644-649bp (AGGCTC/-), strain fgfr4$^{nch5}$ has two 1bp insertion after 605bp (-/G) and 606bp (-/A), and strain fgfr4$^{nch6}$ has one substitution at 609bp (A/T), 38bp deletion at 609-646bp (CAGGAGTATATGTGTGTATGCTACGTGGCACCAAAGAG/-), and a 1bp insertion after 669bp (-/G). Highlighting indicates inserted and mismatched base pairs. (B) Trace sequencing files for all *fgfr4* mutants.

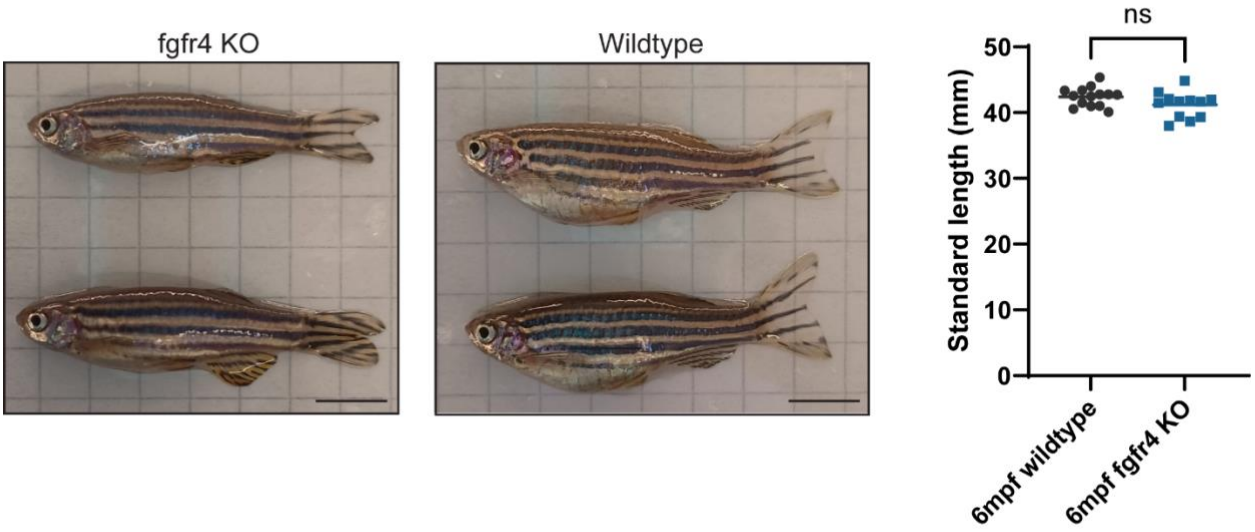


**Supplementary Figure 2. Homozygous *fgfr4* knockout zebrafish are not significantly smaller than wildtype zebrafish at six months post-fertilization.** Standard length quantification was performed with n=14 WT fish and n=12 fgfr4 KO fish. Each point represents an individual fish standard length, and the bar represents the mean. An unpaired two-tailed t-test with Welch's correction was used to calculate the p value. Scale bar is 1 cm.
